# Chromosome Size as a Robust Predictor of Recombination Rate: Insights from Holocentric and Monocentric Systems

**DOI:** 10.1101/2025.07.10.664216

**Authors:** František Zedek, Petr Bureš, Tammy L. Elliott, Marcial Escudero, Kay Lucek, André Marques

## Abstract

Recombination is a fundamental evolutionary process essential for generating genetic diversity, facilitating adaptation, and driving speciation. However, direct measurement of recombination rate remains challenging, as standard methods—such as chiasma counts or genetic linkage maps—are labor-intensive and often infeasible for non-model species. In this study, we identify chromosome number and mean chromosome size as practical proxies for genome-wide recombination rate by analyzing genetic map data from 73 insect species and supplementary analyses of 157 monocentric flowering plants. We confirm the long-standing hypothesis that monocentric species have nearly twice as many crossovers per chromosome as holocentric species, reflecting structural constraints imposed by diffuse centromeres. Using both ordinary and phylogenetically informed Bayesian regression models, we show that recombination rate increases with chromosome number and decreases with mean chromosome size. Crucially, mean chromosome size is a significantly better predictor, particularly in holocentric species. This insight enables recombination rate estimation in thousands of species with known genome and chromosome sizes, thereby allowing hypothesis testing at scales previously unattainable. Building on these results, we present predictive models applicable to poorly studied holocentric plants. Overall, our study highlights the pivotal role of chromosome architecture in recombination evolution and provides an accessible framework for evolutionary genomic research across diverse lineages.

**Article summary:** Why do some species have more genetic shuffling than others? We investigated 73 insect species with detailed genetic maps and found that mean chromosome size is the best predictor of genome-wide recombination rate, not the number of chromosomes. This means we can now estimate recombination rates even in species where genetic maps are unavailable, using only genome size and chromosome number. We also show that chromosome structure matters—species with a single centromere per chromosome (monocentric) have nearly double the recombination compared to those with diffuse centromeres (holocentric). These insights open new doors for studying evolution across many organisms.

## Introduction

The key evolutionary implications of recombination are twofold: recombination can facilitate adaptation across environments by generating and maintaining genetic variation, or it can break up co-adapted allele combinations, potentially disrupting current optimal gene combinations (Ritz et al., 2017; Stapley et al., 2017a). Recombination also plays a crucial role in speciation as the formation of new species often involves a progressive reduction in genetic exchange in specific genomic regions that act as barriers to gene flow (Butlin, 2005). Structural changes along the chromosomes may suppress recombination locally, promoting the evolution of speciation genes (Zhang et al., 2021). On a genome-wide level, a lot of attention has been given to the evolutionary advantage of sex, which promotes recombination. However, the reason why average genome-wide recombination also varies between sexually reproducing organisms is less understood (Stapley et al., 2017a). In sexual eukaryotes, genome-wide recombination rate is determined by the number of crossovers (COs; i.e., chiasmata) per meiosis and by random meiotic segregation of homologs (Stapley et al., 2017b; Otto & Payseur, 2019; Hofstatter et al., 2021). Because estimating recombination rates with standard approaches, such as counting chiasmata or establishing linkage maps (Stapley et al. 2017b), demands time-consuming analysis and is typically impractical for non-model species, it remains challenging to investigate recombination rate-related hypotheses at a macroevolutionary scale across species and environments.

Since at least one CO per chromosome is usually required for proper segregation during meiosis (a phenomenon known as CO assurance; Hunter, 2015) and the formation of additional COs is tightly regulated by CO interference (a mechanism that prevents COs from occurring too closely together; Hunter, 2015), the number of COs per chromosome is maintained within a narrow range (up to 3 COs in 80% of cases; Lian et al., 2023). Therefore, chromosome number is expected to predict genome-wide recombination rate across species (Stapley et al., 2017b). Having more chromosomes also increases the potential for diverse combinations during the assortment of chromosomes in gametes. However, variation in genome size can obscure this relationship.

All else being equal, the necessity of at least one CO per chromosome results in proportionally higher recombination rates in shorter chromosomes (Lynch et al., 2011; Tigano et al., 2022). Heterochromatin is moreover associated with reduced recombination, for instance, in pericentromeric regions, which often accumulate transposable elements and display suppressed CO activity (Kent et al., 2017). Consequently, larger chromosomes, which tend to contain proportionally larger heterochromatic regions (Leitch et al., 2023), may experience a lower rate of recombination per unit length compared to smaller chromosomes. This dynamic could explain the commonly observed negative association between recombination rate and chromosome size of individual chromosomes inside a single karyotype (Kaback et al., 1992; Jensen-Seaman et al., 2004; Backström et al., 2010; Martin et al., 2019; Brazier & Glémin, 2022; Shipilina et al., 2022; Näsvall et al., 2023; Palahí I Torres et al., 2023), which appears to translate to a negative relationship between mean chromosome size and average genome-wide recombination rate (Lynch et al., 2011; Brazier & Glémin, 2022). Thus, chromosome counts and mean chromosome size can potentially be used as proxies for genome-wide recombination rate comparison across a large number of species.

In eukaryotes, two predominant types of chromosomes have evolved that differ regarding their centromere organization, i.e. monocentric and holocentric chromosomes. Importantly, the accuracy with which chromosome number and mean chromosome size predict genome-wide recombination rate is expected to differ between these two types of chromosome organization. For species with monocentric chromosomes, where kinetochores and chromatid cohesion are restricted to a single centromeric region, the number of COs per chromosome mostly varies between one to four (Crismani & Mercier, 2012; Mercier et al., 2015; Blary & Jenczewski, 2019; Otto & Payseur, 2019; Brazier & Glémin, 2022; Lian et al., 2023), but higher numbers have also been reported (Jones & Franklin, 2006; Mancera et al., 2008; Brazier & Glémin, 2022; Lian et al., 2023). In contrast, species with holocentric chromosomes, where kinetochores are maintained along the entire chromosome length (Bureš et al., 2013; Mandrioli & Manicardi, 2020; Hofstatter et al., 2021) cannot rely on the two-step loss of cohesion during meiosis and restrict the CO frequency per homolog to one or two chiasmata at maximum (Schvarzstein et al., 2010; Metlers et al., 2012; Cuacos et al., 2015; Marques & Pedrosa-Harand, 2016; Mandrioli & Manicardi, 2020; Castellani et al., 2024). This upper limit in holocentric species is primarily supported by cytological observations showing that additional chiasmata tend to disrupt proper chromosome segregation (Nokkala et al., 2024; Castellani et al., 2024). The exact mechanism underlying this constraint remains unknown, but it likely relates to the absence of a localized centromere and to the distinct mode of cohesion release along holocentric chromosomes. Such decreased recombination has been shown for both holocentric plants (e.g., Nordenskiöld, 1962; Cabral et al., 2014; Heckmann et al., 2014; Hofstatter et al., 2021; Castellani et al., 2024) and animals (e.g., Barnes et al., 1995; Nokkala et al., 2004; Viera et al., 2009; Lukhtanov et al., 2018). Another difference between holocentric and monocentric species is the composition of chromatin. In holocentric species, chromatin is often not compartmentalized and shows no obvious euchromatic or heterochromatic clusters as in monocentric species (Mandrioli & Manicardi, 2012; Heckmann et al., 2013; Hofstatter et al., 2022; Mata-Sucre et al., 2024) with homogeneous TE distributions along chromosomes (Hofstatter et al. 2022; Shipilina et al., 2022; Mata-Sucre et al., 2024; Castellani et al., 2024). This could potentially result in a more uniform – yet still telomere-biased – distribution of recombination rates along holocentric chromosomes compared to monocentric ones (Haenel et al., 2018; Palahí i Torres et al., 2022; Hofstatter et al., 2022; Shipilina et al., 2022; Castellani et al., 2024). Furthermore, due to their extended kinetochore and the attachment of spindle microtubules along their entire length, holocentric species are more likely to tolerate chromosomal fissions and fusions that have negligible fitness costs (Bureš et al., 2013; Jankowska et al., 2015; Zedek & Bureš, 2019; Zedek et al., 2020; Mandrioli & Manicardi, 2020; Lucek et al., 2022). Therefore, holocentric species are expected to more readily change chromosome numbers and size and, as a consequence, recombination rates in response to selection compared to monocentric species (Bell, 1982; Escudero et al., 2012; Bureš et al., 2013; Escudero et al. 2013; Bureš & Zedek, 2014). Thus, a higher recombination rate in monocentric species is usually expected to be reached by increasing the number of COs per chromosome (Burt, 2000; Stapley et al., 2017b), whereas in holocentric species, a higher recombination rate is predicted to primarily result from increasing chromosome numbers (and thereby decreasing chromosome size).

Although chromosome numbers and mean chromosome size have been used as proxies for genome-wide recombination rate inference in both monocentric (Pessia et al., 2012; Carta et al., 2018) and holocentric species (Bell, 1982; Escudero et al., 2012; Escudero et al., 2013; Spalink et al., 2018; Mackintosh et al., 2019; Márquez-Corro et al., 2021; Elliott et al., 2022), the assumption that these two genomic traits accurately represent recombination rates should be critically assessed. A system encompassing multiple holocentric and monocentric lineages, for which chromosome numbers, genome sizes, and linkage maps or properly counted chiasmata are available to estimate genome-wide recombination rates is required to address these questions. Insects may constitute such a group (Figure 1), where holocentric chromosomes have independently evolved in the orders Lepidoptera, Trichoptera, Hemiptera, Thysanoptera, Psocodea, Zoraptera, Dermaptera, and Odonata, whereas monocentric chromosomes are more broadly found, for example in Diptera, Coleoptera, Hymenoptera, Orthoptera, or Blattodea (Melters et al., 2012; Zedek & Bureš, 2019; Mandrioli & Manicardi, 2020; Marques & Drinnenberg, 2025). Moreover, chromosome number and size, genome size information, and linkage maps are available for a reasonable amount of species of both holocentric and monocentric insect orders.

**Figure 1.**
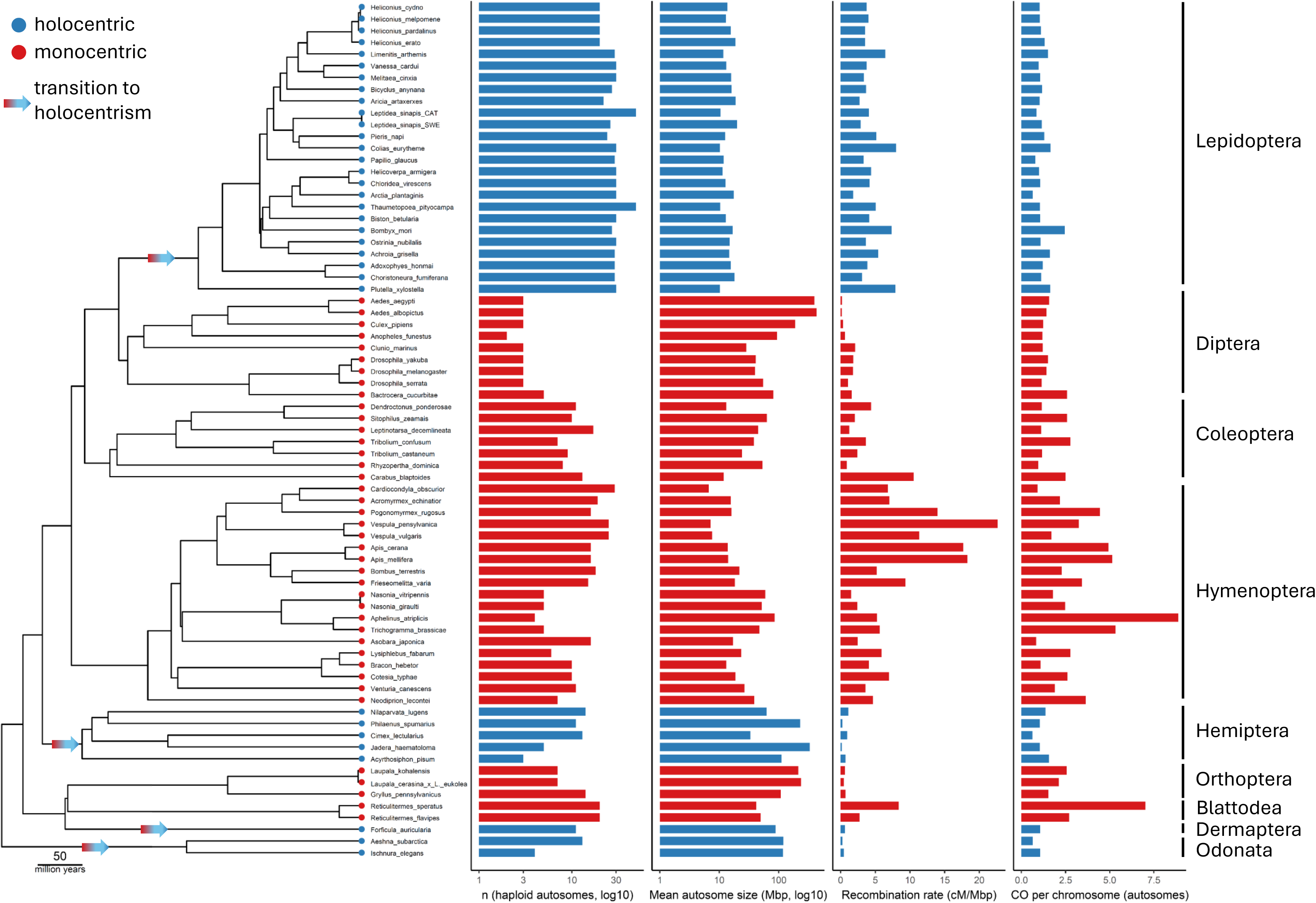
Phylogenetic tree of 33 holocentric (blue) and 40 monocentric (red) species included in our study. Holocentric species include butterflies (Lepidoptera), true bugs (Hemiptera), earwigs (Dermaptera), and dragonflies and damselflies (Odonata). Monocentric species include flies (Diptera), beetles (Coleoptera), sawflies, wasps, bees, and ants (Hymenoptera), termites (Blattodea), and crickets (Orthoptera). The panels with barplots show from right to left haploid autosome numbers (n, log10-transformed), Mean autosome size (Mbp, log10-transformed), autosomal recombination rate (cM/Mb), and number of crossovers per autosome per meiosis. The arrows display putative transitions to holocentric chromosomes (Melters et al., 2021; Marques and Drinnenberg, 2025). The phylogenetic tree is based on the TimeTree resource (Kumar et al., 2022). If a species was not available from TimeTree, we used the tip of its most closely related relative, which was renamed, or added manually based on published phylogenies.

By analyzing recombination data from insect species with well-characterized genetic maps (or counted chiasmata), we aimed to assess the suitability of chromosome numbers and mean chromosome size as predictors of genome-wide recombination rate and test the hypothesis that these predictors are more accurate in holocentric than in monocentric species. Specifically, we expect that (1) the average number of COs per chromosome is higher in monocentric species than in holocentric ones, which is the key assumption stemming from the structural differences between holocentric and monocentric chromosomes (see above), and (2) chromosome number and size predict genome-wide recombination rate better in holocentric than monocentric species. To assess the broader applicability of our results beyond insects, we evaluated the accuracy of chromosome number and mean chromosome size as predictors of recombination rate in flowering plants with monocentric chromosomes. Holocentric species could not be meaningfully included, as only two holocentric plant species currently have established genetic maps (Escudero et al., 2018; Castellani et al., 2024), but we made a prediction of holocentric plants based on the results we obtained from insects and monocentric plants.

## Materials and Methods

### Data Collection

We compiled genetic map length data for 70 species and chiasma count data for three additional species, spanning both holocentric and monocentric insects (Supplementary Table S1). Genome size and chromosome number information were obtained from the NCBI Genome database (Sayers et al., 2022), the Animal Genome Size Database (Gregory, 2025), and additional publications (Supplementary Table S1). We defined the recombination rate as the haploid genetic map length divided by the 1C genome size (Stapley et al., 2017b).

For all but three species, recombination rates were derived directly from available genetic maps. For *Philaenus spumarius* (Hemiptera), *Aeshna subarctica* (Odonata), and *Forficula auricularia* (Dermaptera), where linkage maps were not available, we used published chiasma counts as a proxy for crossover frequency. We converted the mean number of chiasmata per meiosis into a genetic map length by multiplying by 50, assuming that one crossover corresponds to 50 centiMorgans (cM) (Anderson et al., 2004; Chuang & Smith, 2023). We then estimated the mean number of crossovers per chromosome by dividing this value by the haploid chromosome number.

To assess potential artefacts from variation in marker spacing, we tested for associations between map length and average marker distance (cM per marker). No significant correlation was detected (Spearman ρ ≈ 0, p > 0.05) and a linear model controlling for chromosome type likewise showed no effect (β ≈ 0, p > 0.05; Supplementary Figure S1). This confirms that estimates of map length are not biased by marker density.

For all final analyses we used exclusively autosomal map lengths, along with autosomal mean chromosome sizes and autosomal chromosome numbers. This standardization ensures consistency across species, as some published linkage maps report only autosomes. Focusing on autosomes also avoids biases introduced by heteromorphic sex chromosomes, which vary in size among species and are often missing or inconsistently represented in linkage maps. In cases where the size of sex chromosomes (N=14) or the length of sex chromosome maps (N=6) was not reported, we subtracted the average chromosome size or average map length per chromosome from the total genome size or total map length, respectively, to approximate the autosomal component. Genome size estimates were preferentially taken from whole-genome assemblies; this was possible for all but two species, where flow cytometry remained the only source of information. Chromosome counts correspond to the haploid number excluding sex chromosomes, unless explicitly stated otherwise.

### Phylogenetic tree

The phylogenetic tree (Figure 1) used in the following analyses was based on the TimeTree (Kumar et al., 2022), which amalgamates the phylogenetic information of several thousand studies to build an overarching phylogeny of the Eukaryotic Tree of Life. If a species was not available from TimeTree (N=10), we used the tip of its most closely related relative, which was renamed, or added manually based on published phylogenies.

### Recombination rate predicted by chromosome number and mean chromosome size

We analyzed autosomal recombination rate (cM/Mb) and the average number of crossovers (COs) per chromosome in a Bayesian framework using the *brms* R package (Bürkner 2017, 2018, 2021). Because recombination rates are strictly positive and right-skewed, we modeled them on their original scale using a Gamma regression with a log link in all analyses.

We log-transformed the continuous chromosomal predictors and standardized them prior to analysis. Specifically, we considered (i) the haploid number of autosomes, (ii) the mean autosome size (Mb), and (iii) both predictors jointly. Chromosome type (holocentric vs. monocentric) was included as a categorical factor. Models were first specified with interaction terms; if the 95% credible interval for an interaction overlapped with zero, we also analyzed the corresponding additive formulation.

Phylogenetically informed versions of all models were implemented by including a random intercept of the form (1 | gr(tip, cov = A)), where A is the phylogenetic covariance matrix derived from the species tree using the vcv function in the R package *ape* (Paradis & Schliep 2019).

For non-phylogenetic models, we used mildly informative priors: normal(0, 2) on regression coefficients and gamma(2, 0.1) on the Gamma shape parameters. For phylogenetic models, the same priors were applied to coefficients and shape, with an additional exponential(1) prior on the phylogenetic random-effect standard deviation. All models were estimated with four Markov chains and 7–8,000 total iterations per chain, including 2,000 warmup iterations and stringent sampler settings (adapt_delta = 0.99–0.999, max_treedepth = 15).

### Evaluation of the Models and Predictive Accuracy

Model fits were evaluated using approximate leave-one-out cross-validation (LOOIC) and Bayesian R². To further assess predictive accuracy specifically in the comparison between holocentric and monocentric species, we also calculated error metrics based on observed versus fitted recombination rates. We report root mean squared error (RMSE) and mean absolute error (MAE) as absolute measures, and their standardized versions relative to the observed variation in recombination rates (rel_RMSE and rel_MAE). These relative metrics facilitate interpretation by accounting for differences in the magnitude of recombination rates between chromosome types.

### Assessing the relation of crossover number per chromosome to chromosome type

To test whether the average number of COs per chromosome differed between holocentric and monocentric species, we pursued two complementary approaches. Given that the number of COs per chromosome is strictly positive, we first modeled it using a Gamma regression with a log link and used chromosome type as the main predictor. The same priors as in the recombination rate analyses (normal(0,2) for regression coefficients and gamma(2,0.1) for the shape parameter) were applied. Phylogenetic non-independence was incorporated by including a random intercept specified as (1∣gr(phylo, cov = A)).

In addition, we employed Ornstein–Uhlenbeck (OU) phylogenetic modeling to evaluate the evolutionary dynamics of CO number variation between holocentric and monocentric lineages. OU models extend Brownian motion by allowing traits to evolve towards evolutionary optima (θ), thereby testing whether different chromosome types are associated with distinct selective regimes. Analyses were implemented in the R package *OUwie* (Beaulieu et al., 2012). To assign chromosome type states across the phylogeny, we used stochastic character mapping (SIMMAP) under an equal-rates (ER) model, which generates probabilistic reconstructions of ancestral states and thus incorporates uncertainty in the evolutionary history of chromosome type.

We compared seven standard models: two Brownian motion models (BM1 and BMS), which assume a constant variance of the trait through time (BM1 with a single global rate, BMS with regime-specific rates), and five Ornstein–Uhlenbeck models (OU1, OUM, OUMV, OUMA, and OUMVA). OU1 assumes a single optimum θ with constant strength of selection (α) and variance (σ²) across the tree, while OUM allows θ to vary between regimes (holocentric vs. monocentric). OUMV allows both θ and σ² to vary, OUMA allows θ and α to vary, and OUMVA allows all three parameters (θ, α, and σ²) to vary among regimes. Model selection was based on weighted AICs, and the best-fitting model(s) were further evaluated using 500 bootstrap replicates in to obtain empirical estimates and 95% confidence intervals for θ.

### Evaluation of chromosome number and mean chromosome size predictors in angiosperms

To assess the broader applicability of our results beyond insects, we evaluated the accuracy of chromosome number and mean chromosome size as predictors of recombination rate in flowering plants with monocentric chromosomes. Holocentric species could not be meaningfully included, as only two holocentric plant species currently have genetic maps (Escudero et al., 2018; Castellani et al., 2024). To this end, we obtained data on recombination rates, genome sizes, and chromosome numbers from Stapley et al. (2017b) and randomly selected one phylogenetic tree from a set of 100 angiosperm phylogenies provided by Forest (2023). The dataset and phylogenetic tree overlapped with 157 angiosperm species (Supplementary Table S2), which we used in the regression models and their evaluation as described above for insects. To facilitate convergence and comparability of parameter estimates, predictors in phylogenetic models were z-transformed (centered and scaled to unit variance).

## Results

We obtained recombination rate, chromosome number, chromosome size, and the average number of COs per chromosome for 33 holocentric (1 Dermaptera, 5 Hemiptera, 25 Lepidoptera, and 2 Odonata) and 40 monocentric (2 Blattodea, 7 Coleoptera, 9 Diptera, 19 Hymenoptera, and 3 Orthoptera) insect species (Figure 1; Supplementary Table S1).

### Differences in Crossover Number between Chromosome Types

We first assessed whether the average number of COs per chromosome differs between holocentric and monocentric insect species. Using a phylogenetically informed Bayesian Gamma regression, we modeled COs per chromosome as a function of chromosome type with species relatedness included as a random effect (Figure 2A; Supplementary Table S3). The intercept for holocentric species was 0.15 (95% credible interval [CI]: –0.32 to 0.61), corresponding to an expected mean of ≈ 1.16 COs per chromosome. The effect of monocentric species was positive and significant (Estimate = 0.55; 95% CI: 0.02–1.05), implying nearly double the average crossover number relative to holocentric species [≈ 1.82 COs per chromosome].

**Figure 2.**
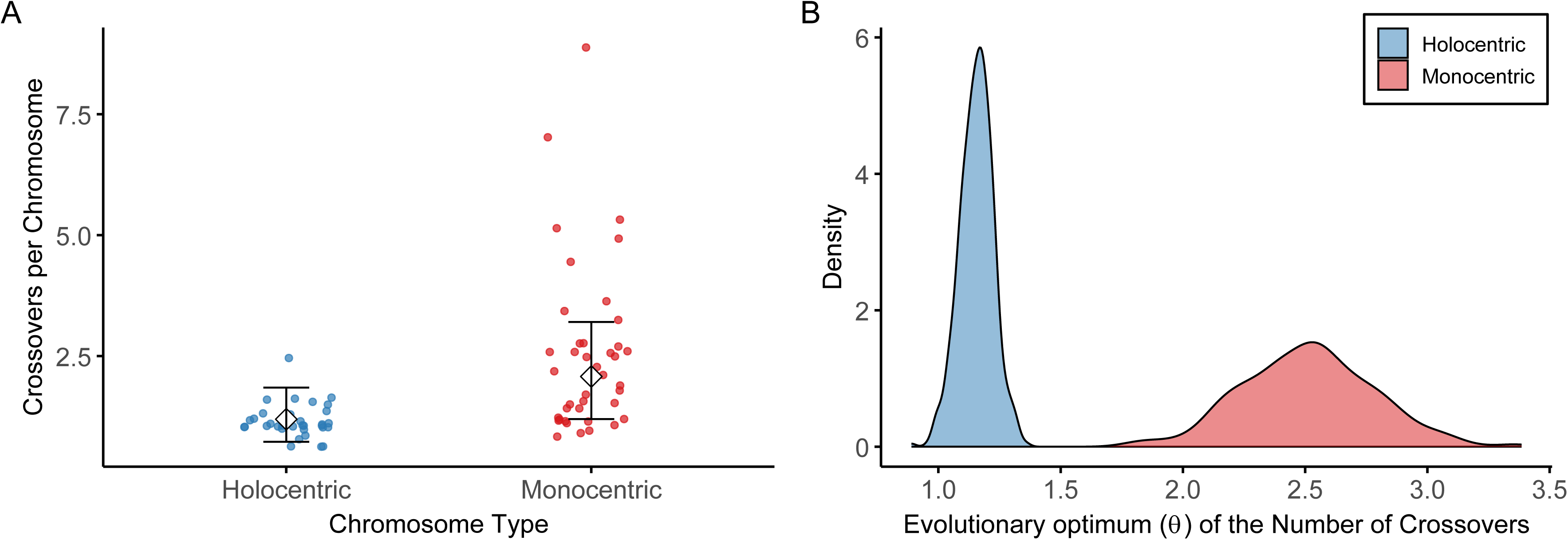
Number of crossovers (COs) in holocentric and monocentric insects. A - Observed COs per chromosome in holocentric (blue) and monocentric (red) species. Individual data points are jittered to display variation, and the overlaid symbols with error bars represent the group mean and 95% prediction credible intervals estimated from the model. B - Bootstrap density distributions (based on 100 replicates) of the evolutionary optimum (θ) for the number of COs estimated under the OUMVA model. The density for holocentric lineages (blue) and monocentric lineages (red) illustrate the uncertainty and differences in the adaptive optima between chromosome types.

To further evaluate the evolutionary dynamics underlying this difference, we applied Ornstein–Uhlenbeck phylogenetic modeling. Across the seven models compared, the OUMV model (allowing optima θ and trait variance σ² to vary between regimes) received overwhelming support (AIC weight ≈ 1.0; Supplementary Table S3). This model estimated distinct adaptive optima for holocentric and monocentric lineages (Figure 2B): θlzl = 1.16 COs per chromosome (95% CI: 1.06–1.23) for holocentric species and θlzl = 2.52 (95% CI: 2.26–2.79) for monocentric species. These results are consistent with the regression analysis and demonstrate that monocentric insects maintain a significantly higher average number of crossovers per chromosome than holocentric insects, in line with expectations from their contrasting meiotic architectures.

### Chromosome Number and Size as Predictors of Recombination Rate

Across the 73 analyzed insect species, recombination rate decreased with increasing mean chromosome size but increased with chromosome number (Figure 3; Supplementary Table S4). Models including mean chromosome size consistently outperformed those including chromosome number in predicting genome-wide recombination rate (Supplementary Table S5). In non-phylogenetic analyses, models with log(mean chromosome size) explained substantially more variance (Bayes R² ≈ 0.61–0.64) than models with chromosome number (Bayes R² ≈ 0.44–0.45). Phylogenetic models confirmed this pattern: those including chromosome size achieved high explanatory power (Bayes R² ≈ 0.85–0.86) with marginal R² values around 0.22–0.29, indicating that chromosome size contributed meaningfully beyond phylogenetic structure (Supplementary Table S5). By contrast, phylogenetic models including chromosome number, despite similar conditional R² values (∼0.92), had marginal R² values near zero (0.02–0.03).

**Figure 3.**
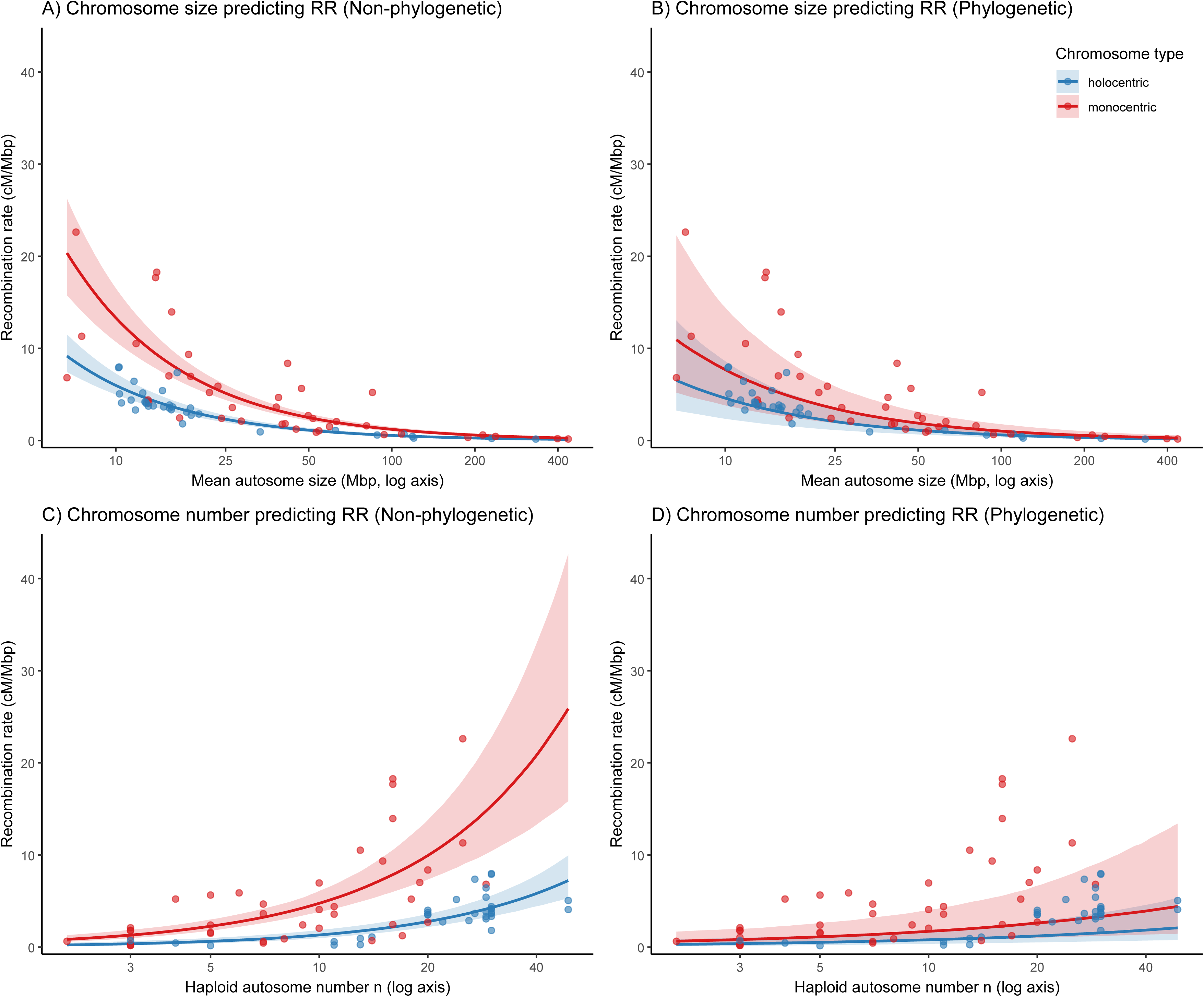
(A) Recombination rate (cM/Mbp) vs. mean chromosome size without accounting for phylogeny. (B) The same relationship when phylogenetic relatedness is considered. (C) Recombination rate vs. haploid autosome number (n) without accounting for phylogeny. (D) Recombination rate vs. haploid autosome number when phylogeny is considered. Solid lines indicate model predictions and shaded ribbons represent 95% prediction credible intervals. Holocentric species are shown in blue, monocentric in red.

Model comparison using leave-one-out cross-validation (LOOIC/ELPD) further highlighted this contrast (Supplementary Table S5). The best-fitting models were consistently those that included chromosome size, while models containing chromosome number performed substantially worse (ΔELPD ≈ –47 to –54). This indicates that the apparent explanatory power of chromosome-number models was almost entirely attributable to phylogenetic signal, whereas chromosome size remained an independent and robust predictor of recombination rate even after accounting for shared ancestry.

For practical use, the best-fitting model (Gamma regression with a log link) can be expressed as:

ln(expected recombination rate) = 4.19 − 1.04 × ln(mean chromosome size [Mbp]) + 0.80 × I(monocentric),

where *I(monocentric)* = 1 for monocentric species and 0 for holocentric species. Here, ln denotes the natural logarithm.

### Assessing Predictive Accuracy in Holocentric and Monocentric Species

In the best-supported non-phylogenetic model (recombination rate ∼ log(mean chromosome size) + chromosome type, see Supplementary Table S5), prediction accuracy differed between chromosome types. Root mean square error (RMSE) and mean absolute error (MAE) quantify the absolute size of prediction errors, while their relative versions (rel_RMSE and rel_MAE) scale these errors by the observed variation in recombination rates. For holocentric species, prediction errors were relatively low (RMSE = 1.13 [95% CI: 1.10–1.36]; MAE = 0.74 [0.67–1.04]; rel_RMSE = 0.52 [0.51–0.63]), indicating a close match between observed and predicted values. In contrast, monocentric species exhibited larger errors (RMSE = 3.89 [3.68–4.79]; MAE = 2.48 [2.38–2.91]; rel_RMSE = 0.73 [0.69–0.89]), suggesting greater unexplained variance. These results demonstrate that recombination rate is more accurately predicted in holocentric species, both in absolute and relative terms. Importantly, the relative metrics show that the poorer fit in monocentrics cannot be explained solely by their higher recombination rates; instead, recombination in monocentric lineages appears intrinsically less predictable from chromosome size.

### Evaluation of Chromosome Number and Mean Chromosome Size as Predictors in Angiosperms

Across 157 angiosperm species with monocentric chromosomes, recombination rate decreased with increasing mean chromosome size but increased with chromosome number (Figure 4; Supplementary Table S6). As in the case of insects, chromosome size was by far the stronger predictor (Supplementary Table S7).

**Figure 4.**
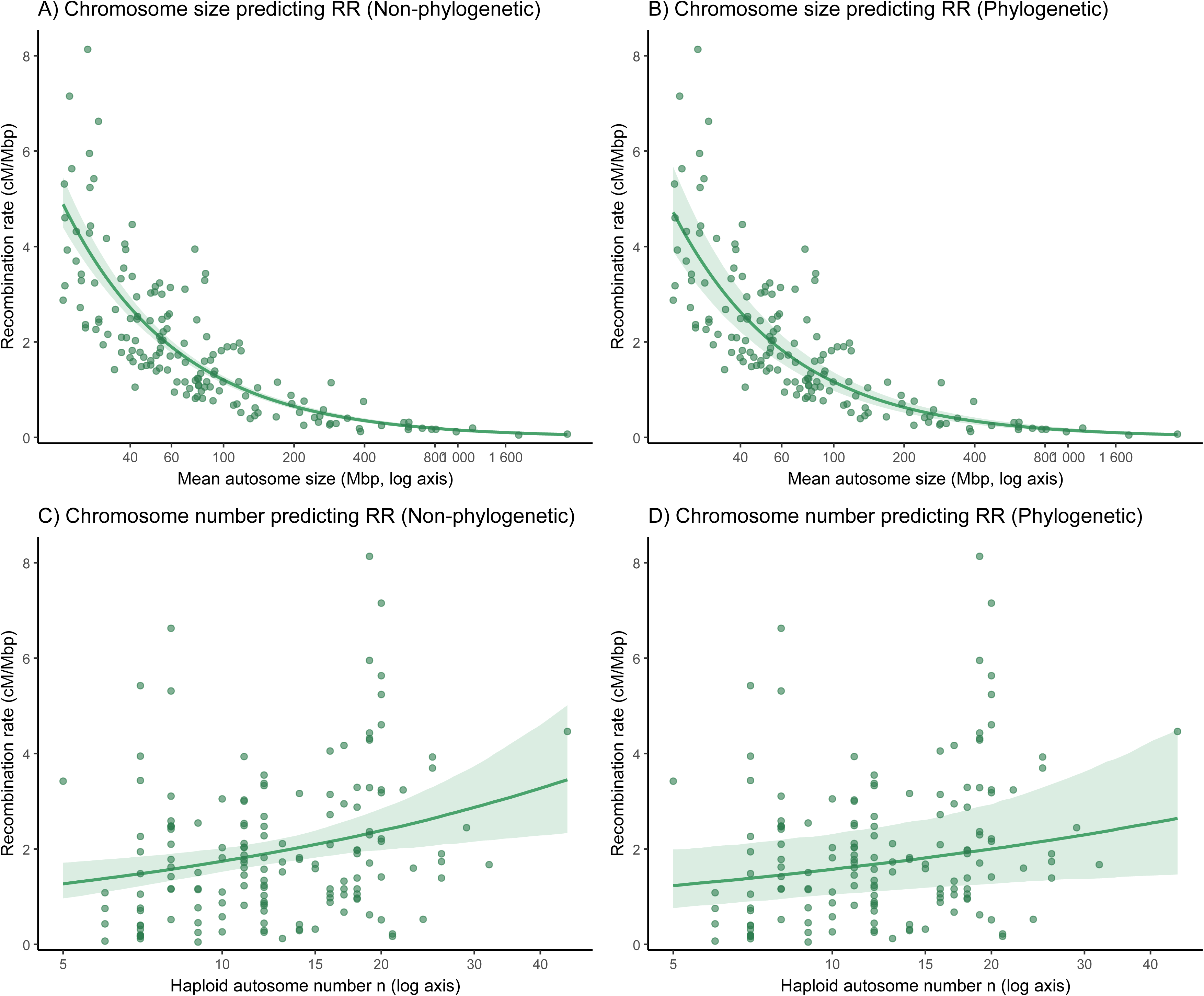
Supplementary Figure S1. Relationships between chromosome number, mean chromosome size, and recombination rate in flowering plants with monocentric chromosomes. (A) Relationship between recombination rate (cM/Mbp) and mean chromosome size. (B) Relationship between recombination rate (cM/Mbp) and mean chromosome size when phylogenetic relatedness is considered. (C) Relationship between recombination rate (cM/Mbp) and haploid chromosome number. (D) Relationship between recombination rate (cM/Mbp) and haploid chromosome number when phylogenetic relatedness is considered. Solid lines indicate model predictions and shaded ribbons represent 95% prediction credible intervals.

In non-phylogenetic analyses, models including log(mean chromosome size) explained substantially more variance (Bayes R² ≈ 0.49–0.50) than those including chromosome number (Bayes R² ≈ 0.07). The estimated effect of size was strongly negative (β ≈ –0.89 ± 0.04), whereas the positive effect of chromosome number was small (β ≈ 0.14 ± 0.06). Phylogenetic models confirmed this pattern: size-based models achieved higher explanatory power (conditional R² ≈ 0.54–0.55; marginal R² ≈ 0.48), while number-based models had much lower values (conditional R² ≈ 0.43; marginal R² ≈ 0.05), indicating that their apparent fit largely reflected phylogenetic structure.

Model comparison using leave-one-out cross-validation further supported these results (Supplementary Table S7). The best-fitting models were consistently those including mean chromosome size, whereas models based on chromosome number performed substantially worse (ΔELPD ≈ –90 to –120). Adding chromosome number or interaction terms did not improve model performance.

Overall, these analyses show that mean chromosome size is a robust, independent predictor of recombination rate in monocentric angiosperms, with smaller chromosomes associated with higher recombination intensity, while the positive relationship with chromosome number is weak and largely phylogenetically driven (Figure 4; Supplementary Tables S6 and S7).

### Predicting recombination in holocentric plants

Building our results from flowering plants (Figure 4), we generated a prediction for holocentric plants by taking the best monocentric plant model (with mean chromosome size as a sole predictor) and shifting its intercept by the monocentric–holocentric offset estimated in insects (Figure 3), while keeping the size slope identical (Figure 5). When we compared this prediction with the two holocentric plants with maps, *Rhynchospora breviuscula* closely matched the predicted curve, whereas *Carex scoparia* lay above it (Figure 5).

**Figure 5.**
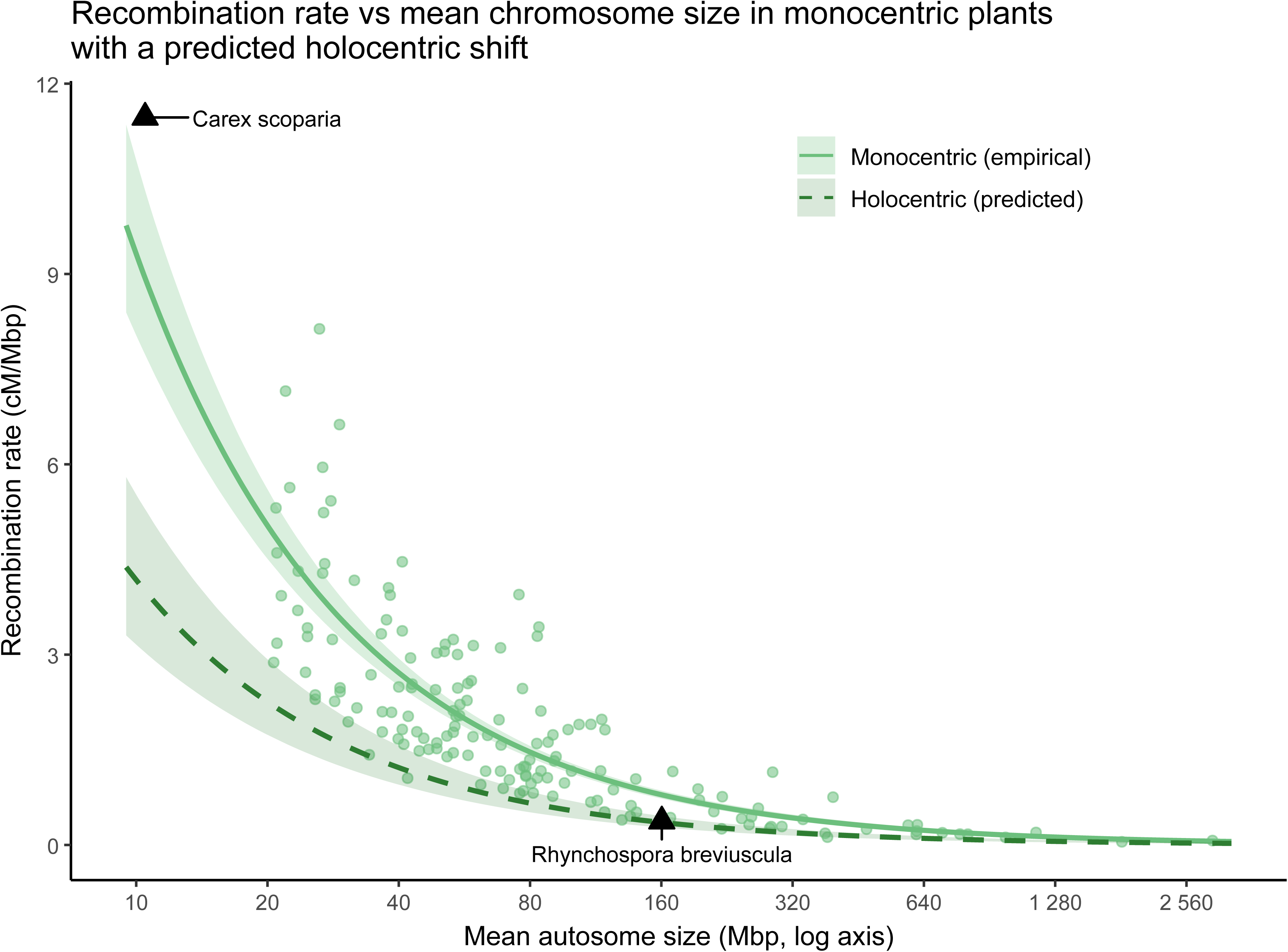
Relationship between mean chromosome size (Mbp) and recombination rate (cM/Mbp) in plants. Light green points represent empirical data for monocentric plants. The light green solid line and shaded area indicate the regression fit and 95% confidence intervals, respectively, for monocentric plants. The dark green dashed line and shaded area represent the predicted regression and 95% confidence intervals for holocentric plants, derived by adjusting the monocentric plant intercept using the intercept difference between monocentric and holocentric insect species. Black triangles show the only two plant species with holocentric chromosomes for which both recombination rate and mean chromosome size are known: *Carex scoparia* (Escudero et al., 2018) and *Rhynchospora breviuscula* (Castellani et al., 2024).

## Discussion

The evolution of recombination rates plays a pivotal role in speciation, balancing the benefits of adaptability with the risks of disrupting established genetic combinations (Ritz et al., 2017; Stapley et al., 2017a). High recombination rates, while fostering evolutionary potential by generating novel allele combinations for adaptation to diverse environments (Wang et al., 2019) or for highly competitive communities (Bell, 1992), can also dismantle favorable co-adapted gene complexes (Ritz et al., 2017). Conversely, lower recombination rates may provide greater reproductive assurance by preserving existing adaptive combinations but at the cost of reduced evolutionary flexibility. This trade-off is particularly evident when considering the factors influencing recombination.

Here, we show that the number of COs per chromosome is predominantly higher and more variable in monocentric insects compared to holocentric ones (Figure 2) which provides support for a long-standing hypothesis based on karyological observations suggesting that holocentric chromosomes typically exhibit more stringent constraints on chiasma formation (White, 1973; Nokkala et al., 2004; Gokhman & Kuznetsova, 2006; Schvarzstein et al., 2010; Metlers et al., 2012; Bureš et al., 2013; Cabral et al., 2014). Mechanistically, these differences likely stem from the centromere organization in holocentric species, which prevents the two-step loss of cohesion typical of monocentric meiosis and often results in only one, or at most two, chiasmata per homologous pair (Schvarzstein et al., 2010; Marques & Pedrosa-Harand, 2016; Hofstatter et al., 2021). However, the exact mechanism underlying this limitation remains unknown. More than two chiasmata per bivalent appear to be either absent or non-viable in holocentric species studied so far (Nokkala et al., 2004; Hollis et al., 2020; Castellani et al., 2024), although the existence of holocentric lineages capable of more than two viable crossovers per chromosome cannot be ruled out. In contrast, monocentric chromosomes can achieve more COs per homolog pair (even over 10) and apparently do so across a range of species (Jones & Franklin, 2006; Fernandes et al., 2018; Otto & Payseur, 2019; Lian et al., 2023).

In line with the tight regulation of CO number per chromosome through CO assurance and interference (Hunter, 2015), both chromosome number and mean chromosome size initially appeared to predict genome-wide recombination rate in insects (Figures 3A,C). However, chromosome number consistently showed lower explanatory power, and across both phylogenetic and non-phylogenetic models, mean chromosome size emerged as the stronger predictor (Figure 3; Supplementary Table S5). Including both variables in additive or interaction models did not improve fit (Supplementary Table S5), indicating that chromosome number contributes little independent information beyond mean chromosome size. This likely reflects that chromosome number captures coarse differences among higher taxonomic groups, while within genera or families, where chromosome numbers vary little (as in Lepidoptera; de Vos et al., 2020), variation in chromosome size more effectively explains subtle differences in recombination rate. Moreover, polyploidization multiplies chromosome number and likely has little effect on genomelzlwide recombination rates (Ross-Ibarra, 2007; Tiley & Burleigh, 2015; Stapley et al., 2017b), although autopolyploids may reduce the CO number per chromosome to avoid multivalent formation (Bomblies et al., 2016). This may further undermine the predictive accuracy of chromosome numbers. In contrast, mean chromosome size effectively controls the multiplicative effects of (neo)polyploidy on chromosome number (Bureš et al., 2024), although there is still unexplained variation left (Figure 3A; Supplementary Table S5). The superior model fit observed for holocentric species across both predictors further suggests that the strict limitations on CO numbers (Figure 2)—imposed by the constraints of their kinetochore architecture—result in more predictable recombination patterns than those seen in monocentric species. Together, these results indicate that mean chromosome size is generally a better proxy for recombination rate and that predictions are more accurate in holocentric than in monocentric systems.

While our findings support that chromosome number and especially mean chromosome size are valid proxies for recombination rate, particularly in holocentric insects, it is important to acknowledge the limitations of our study. Recombination varies along a genome and is undoubtedly influenced by many factors beyond chromosome number and mean chromosome size, such as structural variation, centromere position (metacentric vs. acrocentric in monocentrics), transposable element activity, chromatin structure, and specific meiotic genes (Johnston, 2024). Both sex chromosomes and sex determination systems further differ across our studied insects, which is likely adding additional variation. To estimate the impact of sex determination system and sex chromosomes, data for more species is, however, required with better assemblies for sex chromosomes unique to the heterogametic sex. In addition, the reliability of our approach inevitably depends on the accuracy of genome size and chromosome number estimates. Variation in assembly completeness or sequencing technology can affect genome size estimates and thus the calculated mean chromosome size. These effects are likely minor compared to biological variation but should be considered when applying the model across heterogeneous datasets. Although the broad taxonomic sampling across insects is valuable, extrapolating these findings to other monocentric and holocentric animals — and especially to plants — should be done with caution.

On the other hand, our analyses encompassing 157 monocentric species of flowering plants showed very similar results (Figure 4; Supplementary Tables S6, S7) to our findings in insects (Figure 3, Supplementary Tables S4, S5), suggesting their general applicability. Building on results from insects and monocentric plants, we predicted how genome-wide recombination rate should relate to mean chromosome size in holocentric plants (Figure 5). Notably, *Rhynchospora breviuscula* (Castellani et al., 2024) closely follows the predicted relationship, whereas *Carex scoparia* (Escudero et al., 2018) deviates from it (Figure 5). This deviation may reflect random variation, lineage-specific biological factors, lack of a reference genome assembly, or systematic differences in recombination dynamics between holocentric and monocentric plants. An additional possibility is that *Carex scoparia* lies outside the range of chromosome sizes represented in our monocentric plant dataset; the extrapolated prediction curve may therefore not accurately capture recombination behavior at such small chromosome sizes. Moreover, while our insect analyses suggest a shared slope between chromosome types (Figure 3; Supplementary Table S4), the relationship between chromosome size and recombination rate could differ more fundamentally between monocentric and holocentric plants. However, with data currently limited to just two holocentric plant species, we cannot determine whether the observed deviation is meaningful or incidental. Additional data from diverse holocentric plant lineages will be essential to assess the robustness and generality of our predictions.

Our demonstration of the links among CO number, mean chromosome size, chromosome type, and recombination rate provides a valuable framework for future large-scale evolutionary studies aimed at unraveling the evolution of recombination and its role in adaptation and speciation across the Tree of Life. In light of our findings, it is possible that previous studies on the holocentric plant genus *Carex* (Bell, 1982; Escudero et al., 2012; Escudero et al., 2013; Spalink et al., 2018; Márquez-Corro et al., 2021) and monocentric angiosperms (Carta et al., 2018) may have suffered from reduced power by using chromosome number rather than mean chromosome size as a proxy for recombination rate. However, chromosome numbers are available for many more species because the mean chromosome size requires knowledge of both chromosome number and genome size. For instance, in flowering plants, chromosome numbers are available for ca. 60,000 species (Rice et al., 2015) while mean chromosome size is available for only about 14,000 species (Bureš et al., 2024). Therefore, continuing measurements of genome size (e.g., using flow cytometry) and more initiatives like the Darwin Tree of Life (Darwin Tree of Life Project Consortium, 2022) that generate chromosome-resolved reference genomes for thousands of species will allow us to obtain mean chromosome size data and chromosome numbers at an unprecedented scale.

While holocentric chromosomes appear to offer certain evolutionary advantages, such as the ability to more readily tolerate increased rates of chromosomal fissions and fusions without necessarily reducing fitness (Zedek & Bureš, 2019; Lucek et al., 2022; Zedek et al., 2022)— allowing for rapid karyotypic evolution and potentially increased genetic diversity through altered recombination rates and linkage groups—the prevalence of monocentric chromosomes in the vast majority (80-85%) of eukaryotes (Márquez-Corro et al., 2018) and comparable rates of chromosome number evolution in monocentric and holocentric insects (Ruckman et al., 2020) suggest a different story. This apparent discrepancy–advantages ascribed to holocentrics versus the predominance and comparable dynamics of monocentrics–may be explained by the spatial separation of chromosome segregation and recombination in monocentric chromosomes. In monocentric species, these two crucial meiotic processes are functionally and physically distinct. Holocentric chromosomes, by contrast, must manage both segregation and recombination across their diffuse centromere structure. This constraint likely necessitates a reduction in the number of chiasmata per chromosome to ensure proper segregation, as multiple chiasmata in holocentric species can lead to meiotic errors (Nokkala et al., 2004; Hollis et al., 2020). However, some holocentric insects and plants have evolved non-canonical meioses that decouple where crossovers form from where spindle forces act (Melters et al. 2012; Heckmann et al. 2016; Pedrosa-Harand & Marques 2016; Lukhtanov et al. 2018): inverted (post-reductional) meiosis, in which sister chromatids bi-orient and separate in meiosis I and homologous chromatids segregate in meiosis II, and telokinetic meiosis, which restricts kinetochore activity to one chromosome end so homologs segregate reductionally in meiosis I; additionally, several lineages limit crossovers and remodel meiotic chromosome domains to keep cohesion loss spatially separated from kinetochore function (Pedrosa-Harand & Marques 2016). Our findings, demonstrating a significantly higher number of chiasmata per homolog in monocentric species compared to holocentric species (Figure 2), directly support this explanation. The increased chiasma frequency in monocentrics likely contributes to their greater evolutionary success by allowing for higher levels of genetic recombination. Thus, while holocentric species may offer flexibility in karyotypic evolution, the inherent trade-off between recombination rate specifically for chiasma formation and proper segregation, might ultimately limit their overall evolutionary success compared to the more specialized and efficient mechanisms found in monocentric chromosomes. In summary, our findings reveal a fundamental difference in how insect recombination is regulated. While monocentric species rely on increased CO rates, holocentric species appear to compensate through changes in chromosome number. This stark contrast underscores centromere architecture’s critical role in shaping insect genomes’ evolutionary trajectory.

## Data availability statement

All the data used in this article are available in supplementary materials online on the publisher’s website.

## Author contributions

FZ conceived the study and performed analyses. FZ, PB, TLE, ME, KL, and AM wrote the manuscript.

---

**Supplementary Figure S1.**
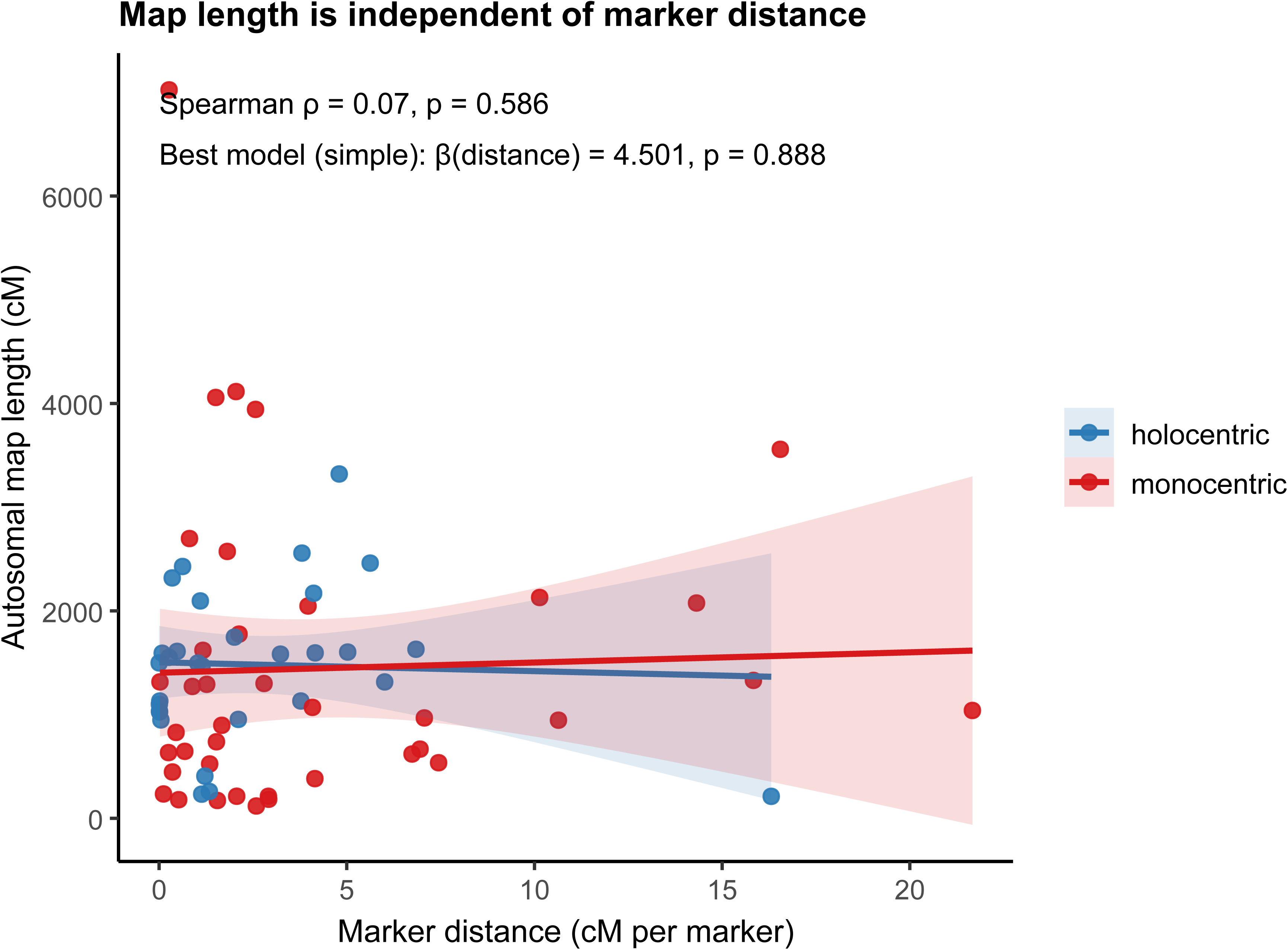
Associations between genetic map length and average marker distance (cM per marker). No significant correlation was detected (Spearman ρ ≈ 0, p > 0.05) and a linear model controlling for chromosome type likewise showed no effect (β ≈ 0, p > 0.05). This confirms that estimates of map length are not biased by marker density.

